# Generalizing movement patterns following shoulder fixation

**DOI:** 10.1101/678623

**Authors:** Rodrigo S. Maeda, Julia M. Zdybal, Paul L. Gribble, J. Andrew Pruszynski

## Abstract

A common goal of motor learning is generalizing newly learned movement patterns beyond the training context. Here we tested whether learning a new physical property of the arm during self-initiated reaching generalizes to new arm configurations. One hundred human participants performed a single-joint elbow reaching task and/or countered mechanical perturbations that created pure elbow motion. Participants did so with the shoulder joint either free to rotate or locked by the robotic manipulandum. With the shoulder free, we found activation of shoulder extensor muscles for pure elbow extension trials, as required to counter the interaction torques that arise at the shoulder due to forearm rotation. After locking the shoulder joint, we found a substantial reduction in shoulder muscle activity that developed slowly over many trials. This reduction is appropriate because locking the shoulder joint cancels the interaction torques that arise at the shoulder to do forearm rotation and thus removes the need to activate shoulder muscles. In our first three experiments, we tested whether this reduction generalizes when reaching is self-initiated in (1) a different initial shoulder orientation, (2) a different initial elbow orientation and (3) for a different reach distance/speed. We found reliable generalization across initial shoulder orientation and reach distance/speed but not for initial elbow orientation. In our fourth experiment, we tested whether generalization is also transferred to feedback control by applying mechanical perturbations and observing reflex responses in a distinct shoulder orientation. We found robust transfer to feedback control.

**New & Noteworthy:** Here we show that learning to reduce shoulder muscles activity following shoulder fixation generalizes to other movement conditions but does not generalize globally, indicating that the nervous system does not implement such learning by modifying a general internal model of arm dynamics.

**Disclosures:** The authors declare no conflict of interest.

## Introduction

The nervous system must coordinate muscles across many joints when generating motor actions. For example, performing a single joint elbow movement requires contracting muscles at the shoulder joint to compensate for rotational forces that arise at the shoulder during forearm rotation (Ghez and Sainburg 1995; Hollerbach and Flash 1982; Virji-Babul and Cooke 1995). The predictive nature of this shoulder contraction, and the fact that it scales appropriately with the required shoulder torque as a function of planned movement speed and initial arm configuration, suggests that the control signals sent to arm muscles rely on an internal model of the arm’s intersegmental dynamics (Gribble and Ostry 1999; Maeda et al. 2017).

We have recently showed that people slowly reduce shoulder muscle contraction when generating pure elbow movements after mechanically fixing the shoulder joint (Maeda et al. 2018). Learning to reduce shoulder muscle activity is efficient in this scenario because mechanically fixing the shoulder joint cancels the interaction torques that arise at the shoulder when the forearm rotates and thus negates the need to recruit shoulder muscles. Here we begin to address how this learning arises by investigating whether, and to what extent, learning generalizes to other experimental conditions - a well-established approach for gaining insight into the structure of motor learning (for review, see Krakauer et al. 2019; Shadmehr 2004). Given that the predictive shoulder contraction itself likely relies on an internal model of the arm’s dynamics, one possibility is that learning to reduce shoulder muscle activity after fixing the shoulder joint arises because the nervous system is updating an internal model of the arm. If so, then this learning should generalize broadly to movements at different speeds, distances and with different arm configurations. Another option is that the nervous system may treat shoulder fixation as a local feature within a new environment and, consistent with many studies investigating motor learning in such scenarios, exhibit limited generalization to movements at different speeds and different arm configurations (Berniker and Kording 2008; Brayanov et al. 2012; Burgess et al. 2007; Criscimagna-Hemminger et al. 2003; Ingram et al. 2010; Krakauer et al. 2000; Malfait et al. 2002, 2005; Sainburg et al. 1999; Shadmehr and Moussavi 2000). Lastly, the nervous system may not generalize at all because, unlike learning a new force field or visuomotor environment, the intervention used in this paradigm does not cause participants do not make substantial or systematic kinematic errors.

As in our previous work (Maeda et al. 2018), participants performed single-joint elbow movements with the shoulder free to rotate or with the shoulder locked by the robotic manipulandum. We then tested whether learning these altered arm dynamics during reaching movements generalized to self-initiated reaching (i) in a different initial shoulder orientation, (ii) in a different initial elbow orientation and (iii) for a different reach distance/speed in the same initial shoulder and elbow orientations. We found reliable generalization across initial shoulder orientation and reach distance/speed but not for initial elbow orientation. We also tested whether learning these altered arm dynamics during reaching generalizes to feedback responses (i.e. stretch reflexes) in a different shoulder orientation and found reliable generalization. Taken together, our results show that learning to reduce shoulder muscles activity following shoulder fixation does generalize to other movement conditions but that such generalization is not universal, suggesting that the nervous system does not implement such learning by modifying its internal model of their arm’s dynamics.

## Materials and Methods

### Subjects

A total of 100 healthy volunteers (aged 17–47, 57 females) participated in one of four experiments. All participants reported that they were right-handed and had no history of visual, neurological, or musculoskeletal disease. Participants provided written informed consent, were free to withdraw from the experiment at any time, and were paid for their participation. The Office of Research Ethics at Western University approved this study.

### Experimental task and apparatus

All experiments were performed using a robotic exoskeleton (KINARM, Kingston, ON, Canada) that permits flexion and extension movement of the shoulder and elbow joints in the horizontal plane and can selectively apply torque at both joints (Pruszynski et al. 2008, 2009; Scott 1999). Targets and calibrated hand aligned cursor feedback were projected into the horizontal plane of the task via an LCD monitor and a semi-silvered mirror. Direct vision of the arm was prevented with a physical shield. The two segments of the exoskeleton robot (upper arm and forearm) were adjusted to each participant’s arm and were filled with high-density foam to ensure tight coupling with the robot’s links.

In all experiments, participants began a trial by moving the projected hand cursor to a home target (red circle; 0.6-cm diameter). We displayed the home target so that the elbow and shoulder joints were positioned at the initial orientation required for each experiment (Figure 1, top left). After remaining at this location for a random period of time (250-500 ms, uniform distribution), a goal target was presented in a location that could be reached with a pure elbow extension movement. After another random period (250-500 ms, uniform distribution), the goal target turned red, the hand feedback cursor was extinguished and participants were allowed to start the movement. The hand feedback cursor remained off for the duration of the movement. Participants were instructed to move quickly to the goal target with a movement time between 100 and 180 ms (computed as the time between exiting the home target to entering the goal target). If movement was faster than 100 ms the goal target turned orange, if it was slower than 180 ms the target turned red, otherwise it turned green. No restrictions were placed on movement trajectories. After achieving the goal target, participants were instructed to remain at that location for an additional 500 ms to finish a trial. After a random period (0-1 s, uniform distribution), the goal target became the new home target (0.6 cm diameter) for a flexion movement and the same procedures were repeated.

**Figure 1:**
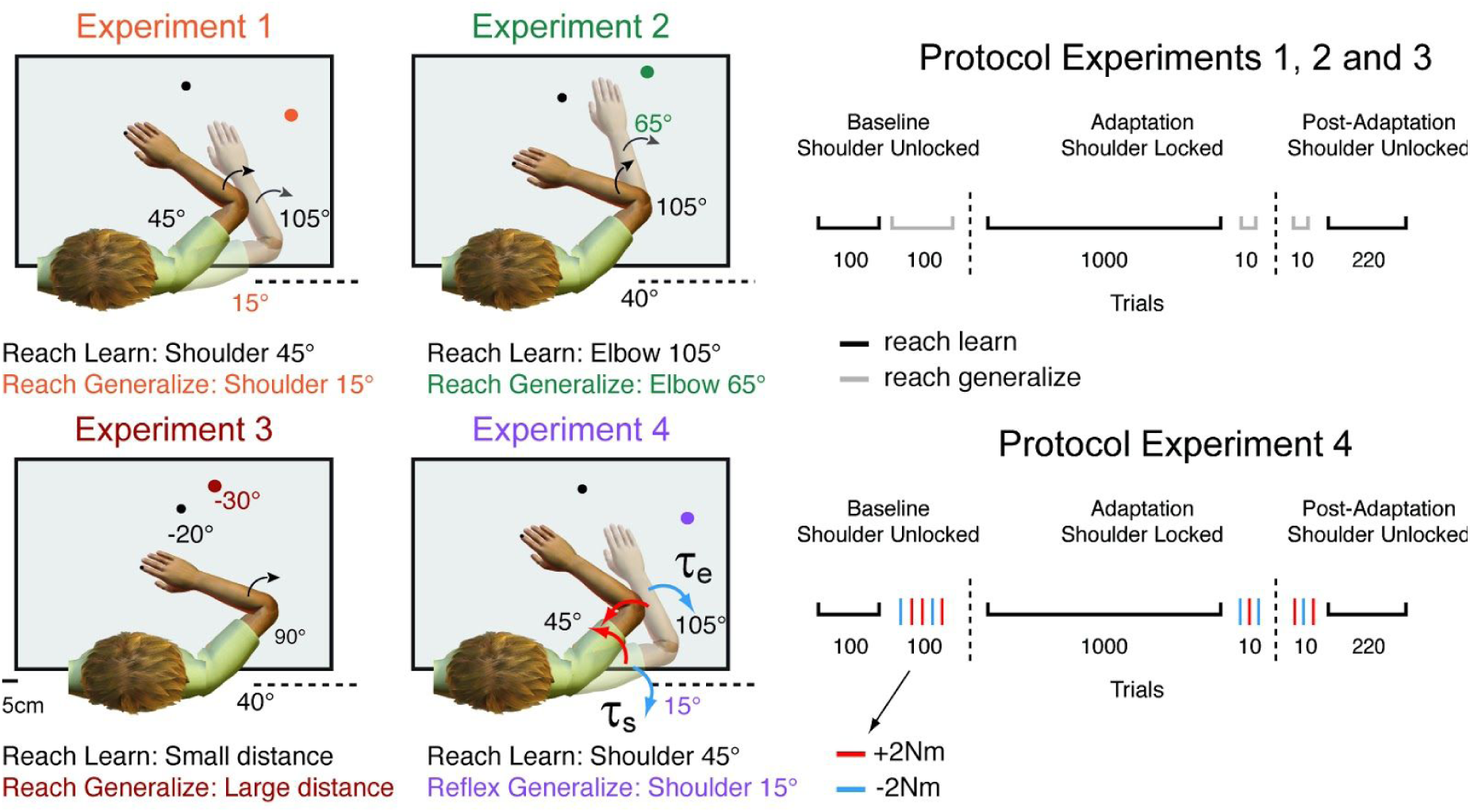
Experimental setup. In Experiments 1, 2, 3 and 4, participants were presented with a peripheral target that could be achieved by rotating only the elbow joint. Participants performed the same movement with the shoulder joint free to rotate (normal arm dynamics) and with the shoulder joint locked by the robotic manipulandum (altered arm dynamics)—reach learn. In Experiments 1, 2 and 3, participants performed a few reaching trials (reach generalize) before and after learning to probe generalization of this learning when reaching with a distinct initial shoulder orientation, with a distinct initial elbow orientation, and when reaching to a distinct reach distance/speed (left panels—experiments 1, 2, and 3). In Experiment 4, mechanical perturbations were applied (reflex generalize) to test the sensitivity of reflex responses before and after learning in a distinct initial shoulder orientation (left panel—Experiment 4). Red and blue arrows represent the direction of the multi-joint step-torques applied to the shoulder and elbow joints. Illustrations of the protocols for Experiments 1, 2 and 3 are shown on the top right panel. Illustration of the protocol for Experiment 4 is shown on the bottom right panel. In Experiments 1, 2, 3 and 4, participants performed 100 extension and flexion baseline trials with the shoulder joint unlocked in the learning condition (black marks) and 100 baseline trials in the generalization condition (reach generalize – gray marks or reflex generalize – red and blue tick marks), 1000 adaptation trials with the shoulder joint locked in the learning condition, 10 adaptation trials in the generalization condition, and 10 post-adaptation trials with the shoulder joint unlocked in the generalization condition, followed by 220 post-adaptation trials in the learning condition.

In selected trials of Experiment 4, when the hand cursor entered the home target, the exoskeleton gradually applied (over 2 s) a background torque of (−2 Nm/+2Nm) to the elbow to ensure baseline activation of shoulder and elbow muscles. After maintaining the cursor in the home target for a randomized duration (1.0–2.5 s, uniform distribution), a step-torque (i.e., perturbation) was applied to the shoulder and elbow joints (2 Nm at each joint over and above the background torque, 100ms duration), which displaced the participant’s hand outside the home target. We chose this combination of shoulder and elbow loads to minimize shoulder motion (see Kurtzer et al. 2014, 2008; Kurtzer 2019; Maeda et al. 2017, 2018). Participants were instructed to quickly counter the load and bring their hand back to the goal target (centered on the home target). If the participant returned to the goal target within 385 ms of perturbation onset, the target circle changed from white to green, otherwise the target circle changed from white to red. In five percent of all trials, the background torques turned on, remained on for the same time period (1.0–2.5 s, uniform distribution), but then slowly turned off, after which participants were still required to perform the reaching movements. These trials ensured that background loads were not fully predictive of perturbation trials.

### Experiment 1: Generalization to a different shoulder configuration

Fifteen participants performed twenty-five degree elbow extension and flexion movements with the shoulder joint free to move and with the shoulder fixed starting from two different initial shoulder orientations (45° for learning trials (reach learn) and 15° for trials in which we probed for generalization (reach generalize).

Participants first completed 100 twenty-five degree elbow extension and flexion trials with the initial shoulder orientation at 45° and 100 twenty-five degree elbow extension and flexion trials with the initial shoulder orientation at 15°, with the shoulder joint free to move (baseline trials). We then mechanically locked the shoulder joint of the KINARM with a physical clamp and participants completed 1000 twenty-five degree elbow extension and flexion trials with the initial shoulder orientation at 45° degrees (adaptation phase in the reach learn condition). At the end of this phase, participants generated 10 twenty-five degree elbow extension and flexion trials with the initial shoulder orientation at 15° degrees but with the shoulder locked at this new orientation (generalization phase). We then unlocked the shoulder joint and participants again generated the same twenty-five degree of elbow extension and flexion movements with the initial shoulder orientation at 15° degree for an additional 10 trials (post-adaptation phase). Lastly, participants again generated twenty-five degree elbow extension and flexion movements for an additional 110 trials with the shoulder initial orientation at 45° degrees (post-adaptation phase) (Figure 1, top right column).

To ensure that all the participants returned to baseline levels with the initial shoulder orientation at 15°, participants also performed an additional 20 trials in that condition but this data was not analyzed given that this washout happened quickly consistent with the washout in the learning posture (Maeda et al. 2018).

Experiment 1 took about 2.5 hours. Rest breaks were given when requested. Prior to data collection, participants completed practice trials until they achieved ∼80% success rates (approx. 5 min).

### Experiment 2: Generalization to a different elbow configuration

Fifteen participants performed twenty-five degree elbow extension and flexion movements with the shoulder joint free to move and with the shoulder fixed starting from two different initial elbow orientations (105°, reach learn condition and 65°, reach generalize condition).

Participants first completed 100 twenty-five degree elbow extension and flexion trials with the initial elbow orientation at 105° and 100 twenty-five degree elbow extension and flexion trials with the initial elbow orientation at 65°, with the shoulder joint free to move (baseline trials). We then mechanically locked the shoulder joint of the KINARM with a physical clamp and participants completed 1000 trials of twenty-five degree elbow extensions and flexions with the initial elbow orientation at 105° degrees (adaptation phase). At the end of this phase, participants generated 10 twenty-five degree elbow extension and flexion movements with the initial elbow orientation at 65° (generalization phase). We then unlocked the shoulder joint and participants again generated the same twenty-five degree elbow extension and flexion movements with the initial elbow orientation at 65° for an additional 10 trials (post-adaptation phase). Lastly, participants again generated twenty-five degree elbow extension and flexion movements for an additional 110 trials with the initial elbow orientation at 105° degrees (post-adaptation phase) (Figure 1, top right column).

Again, to ensure that all the participants returned to baseline levels with the initial elbow orientation at 65°, participants also performed an additional 20 trials in that condition but this data was not analyzed given that this washout took place quickly consistent with previous work (Maeda et al. 2018).

Experiment 2 took about 2.5 hours. Rest breaks were given when requested. Prior to data collection participants completed practice trials until they achieved ∼80% success rates (approx. 5 min).

### Experiment 3: Generalization to a different reach distance/speed

Fifteen participants performed twenty-degree (reach learn) and thirty-degree (reach generalize) elbow extension and flexion movements with the shoulder joint free to move and with the shoulder fixed.

Participants first completed 100 twenty-degree elbow extension and flexion trials and 100 thirty-degree elbow extension and flexion trials, with the shoulder joint free to move (baseline trials). We then mechanically locked the shoulder joint with a physical clamp and participants completed 1000 trials of twenty-degree elbow extensions and flexions (adaptation phase). At the end of this phase, participants generated 10 thirty-degree elbow extension and flexion movements (generalization phase). We then unlocked the shoulder joint and participants again generated 10 thirty-degree elbow extension and flexion movements (post-adaptation phase). Lastly, participants generated twenty-degree elbow extension and flexion movements for an additional 110 trials (post-adaptation phase) (Figure 1, top right column).

Participants also performed an additional 20 trials of thirty-degree elbow extension and flexion movements to ensure that participants returned to baseline levels in this condition. This data was not analyzed given that this washout took place quickly (Maeda et al. 2018).

Experiment 3 lasted about 2.5 hours. Rest breaks were given when requested. Prior to data collection participants completed practice trials until they achieved ∼80% success rates (approx. 5 min).

### Experiment 4: Generalization to feedback responses in a different shoulder configuration

Fifteen participants performed twenty-degree elbow extension and flexion movements with the shoulder joint free to move and with the shoulder fixed (reach learn, shoulder joint at 45°), and countered mechanical perturbations that caused pure elbow motion with the shoulder joint in a different initial shoulder orientation (reflex generalize, shoulder joint at 15°).

Participants first completed 100 twenty-degree elbow extension and flexion trials, and countered mechanical perturbations with the initial shoulder orientation at 45°, with the shoulder joint free to move (baseline trials). In addition, participants performed 100 twenty-degree elbow extension and flexion trials, and countered mechanical perturbations with the initial shoulder orientation at 15°, with the shoulder joint free to move (baseline trials). We then mechanically locked the shoulder joint with a physical clamp and participants generated 1000 twenty-degree elbow extension and flexion trials with the initial shoulder orientation at 45° degree (adaptation phase). At the end of this phase, participants compensated for mechanical elbow perturbations (10 trials) with the initial shoulder orientation at 15° (generalization phase). We then unlocked the shoulder joint and participants compensated for shoulder and elbow perturbations that created pure elbow motion (10 trials) (post-adaptation phase). Lastly, participants again generated twenty-degree elbow extension and flexion movements for an additional 110 trials with the initial shoulder orientation at 45° degree (post-adaptation phase). (Figure 1, bottom right column).

Participants also compensated for mechanical elbow perturbations (20 trials) with the initial shoulder orientation at 15° to ensure that participants returned to baseline levels in this condition. This data was not analyzed given that this washout took place quickly (Maeda et al. 2018).

The order of all perturbation, control and reaching trials was randomized in the baseline and post-adaptation phases and randomized in blocks early and late in the adaptation phase. Experiment 4 lasted about 2.5h. Rest breaks were given throughout or when requested. Prior to data collection participants completed practice trials until they comfortably ∼80% success rates (approx. 5 min).

### Control Experiments

We performed two control experiments. First, ten additional participants performed the same version of Experiments 1 and 4 without locking the shoulder joint for the 1000 trials that would have made up the adaptation phase. This served as a control to rule out changes in feedforward or feedback control caused by extensive practice rather than the shoulder locking manipulation. Second, thirty additional participants also performed the learning phases (baseline with shoulder joint unlocked, adaptation with shoulder locked and post-adaptation with shoulder unlocked) in the generalization conditions of Experiments 1 (N = 10), 2 (N = 10) and 3 (N = 10). This served as a control to ensure that learning to reduce shoulder muscle activity takes place independent of the different conditions used when probing for generalization.

### Kinematic recordings and analysis

Movement kinematics (i.e. hand position and joint angles) were sampled at 1000 Hz and then low-pass filtered (12 Hz, 2-pass, 4th-order Butterworth). In Experiments 1, 2 and 3, all data were aligned on movement onset. In Experiment 4, data from reaching trials was aligned on movement onset and data from perturbation trials was aligned on perturbation onset. Movement onset was defined as 5% of peak angular velocity of the elbow joint (see Gribble and Ostry 1999; Maeda et al. 2017, 2018). We quantified the adaptation and aftereffects of reaching movements following shoulder fixation using hand path errors relative to the center of the target at 80% of the movement between movement onset and offset (the latter also defined at 5% of peak angular velocity of the elbow joint). On average, 80% of the movement corresponds to 170 ms (SD 15 ms) after movement onset. This moment was chosen to capture the state of the kinematics before any feedback corrections. For Experiments 1, 2 and 3, we used software provided by BKIN Technologies in MATLAB to calculate shoulder and elbow torques (Figures 2-4, inset) based on anthropometric values for each participant and a model of the KINARM robot linkages.

**Figure 2:**
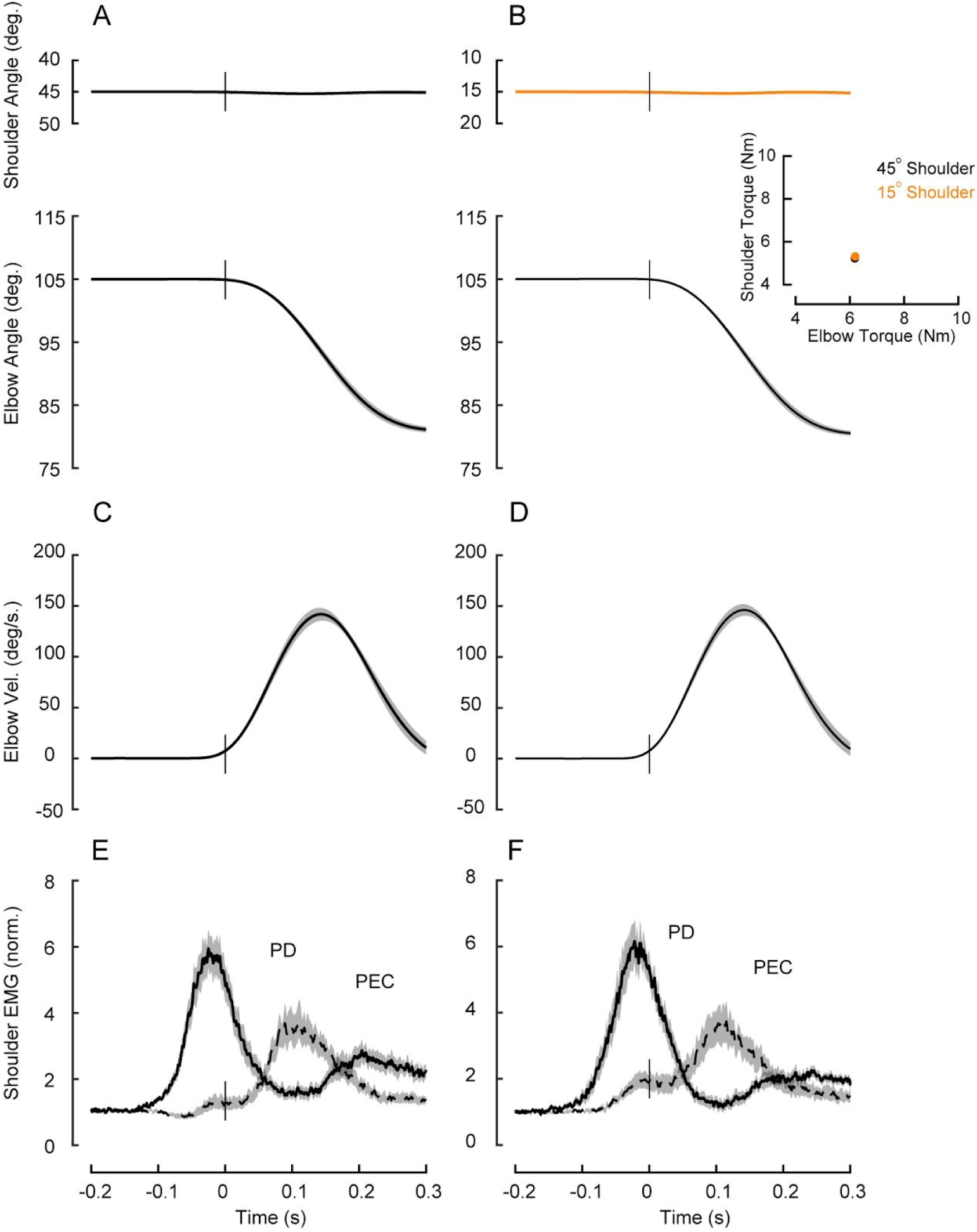
Baseline reaching behaviour in Experiment 1. **A-B**, Average shoulder and elbow angle of elbow extension trials starting from 2 different initial shoulder orientations. Shaded error areas represent the standard error of the mean (SE). Data are aligned on movement onset. Inset shows peak shoulder torque by peak elbow torque separate for each shoulder initial orientation condition. **C-D**, Data of elbow velocity for the elbow kinematic profiles in **A-B**, respectively. Shaded error areas represent SE. Data are aligned on movement onset. **E-F**, Solid and dashed lines represent average agonist (PD) and antagonist (PEC) muscle activity during elbow extension movements for the kinematic profiles in **A-B**, respectively. EMG normalized as described in the Methods. Data aligned on movement onset. Shaded error areas represent the SE.

**Figure 3:**
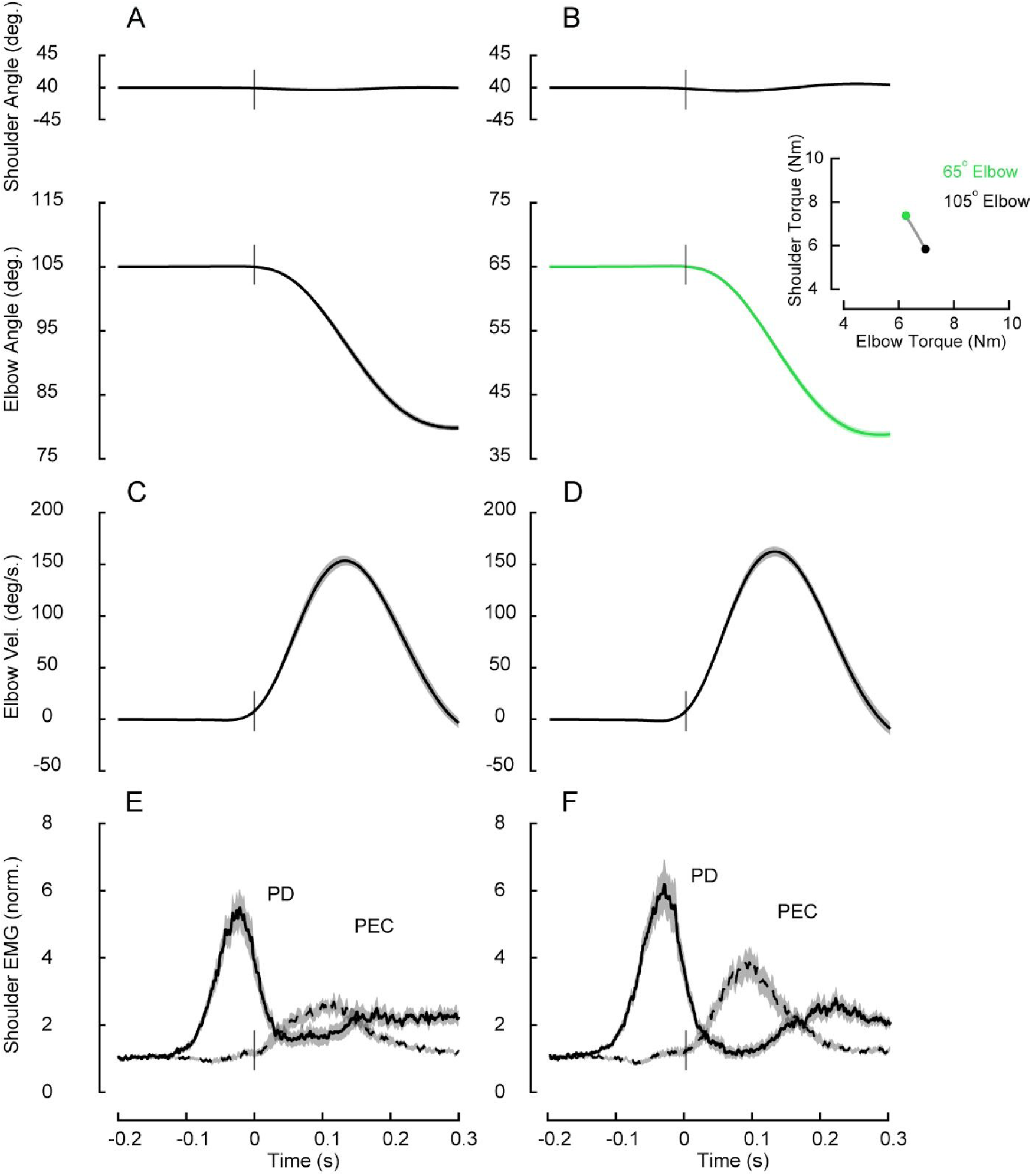
Baseline reaching behaviour in Experiment 2. **A-B**, Average shoulder and elbow angle of elbow extension trials starting from 2 different elbow initial orientations. Shaded error areas represent the standard error of the mean (SE). Data are aligned on movement onset. Inset shows the peak shoulder torque by peak elbow torque separate for each elbow initial orientation. **C-D**, Data of elbow velocity for the elbow kinematic profiles in **A-B**, respectively. Shaded error areas represent SE. Data are aligned on movement onset. **E-F**, Solid and dashed lines represent average agonist (PD) and antagonist (PEC) muscle activity during elbow extension movements for the kinematic profiles in **A-B**, respectively. EMG normalized as described in the Methods. Data aligned on movement onset. Shaded error areas represent the SE.

**Figure 4:**
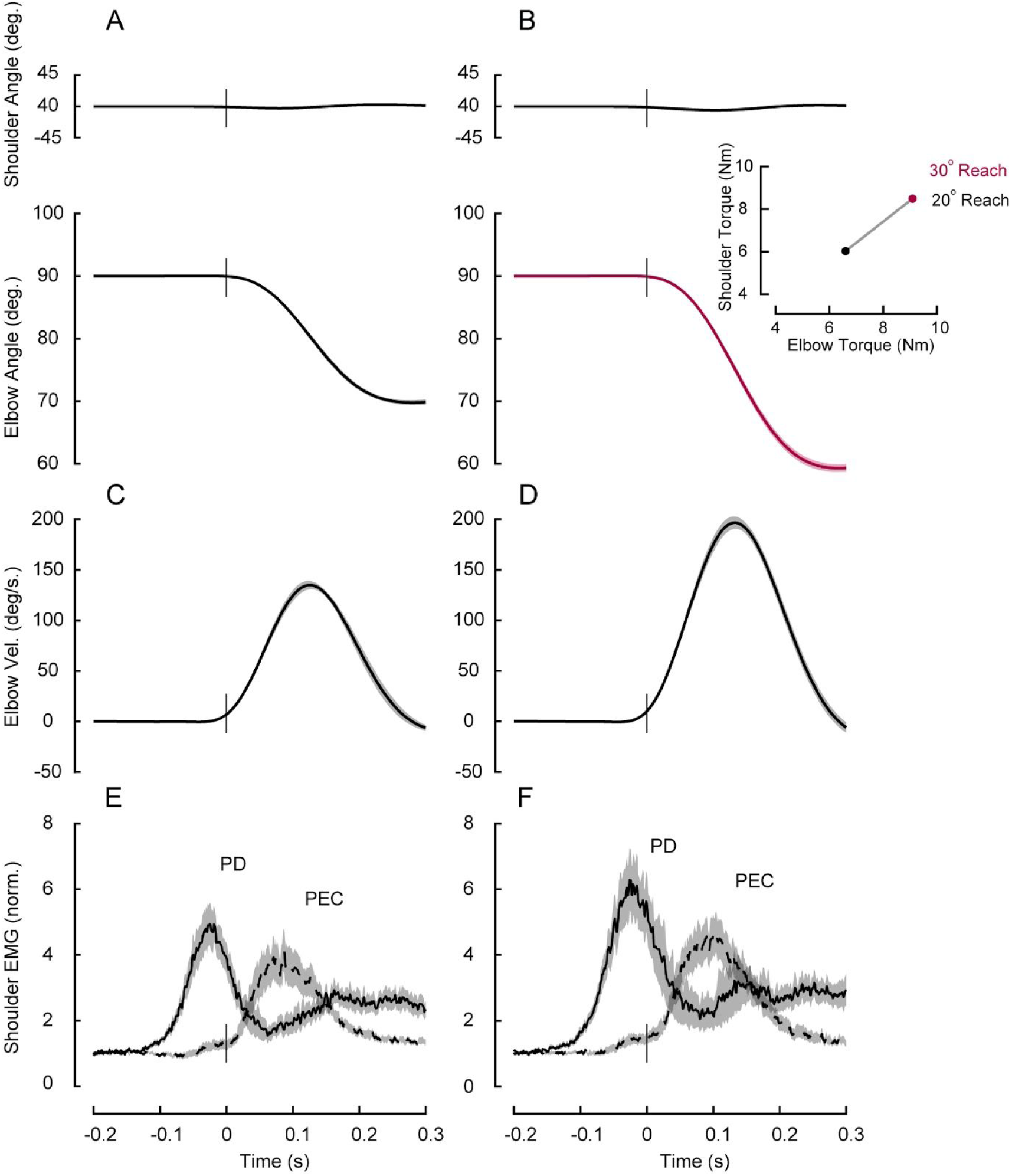
Baseline reaching behaviour in Experiment 3. **A-B**, Average shoulder and elbow angles for 20 degree and 30 degree movements of elbow extension trials, respectively. Shaded error areas represent the standard error of the mean (SE). Data are aligned on movement onset. Inset shows the calculated peak shoulder torques by peak elbow torques for each reach condition. **C-D**, Data of elbow velocity for the elbow kinematic profiles in **A-B**, respectively. Shaded error areas represent the SE. Data are aligned on movement onset. **E-F**, Solid and dashed lines represent average agonist (PD) and antagonist (PEC) muscle activity during elbow extension movements for the kinematic profiles in **A-B**, respectively. EMG normalized as described in the Methods. Data aligned on movement onset. Shaded error areas represent the standard error (SE).

### EMG recordings and analysis

We used commercially available surface electrodes and amplifiers (Delsys Bagnoli-8 system with DE-2.1 sensors, Boston, MA) to measure electromyographic signals from upper limb muscles. Electrodes were placed on the muscle belly and in parallel to the orientation of muscle fibers of five muscles (pectoralis major clavicular head, PEC, shoulder flexor; posterior deltoid, PD, shoulder extensor; biceps brachii long head, BB, shoulder and elbow flexor, Brachioradialis, BR, elbow flexor; triceps brachii lateral head, TR, elbow extensor). Before placing the electrodes, the skin was abraded with rubbing alcohol, and the electrodes were coated with conductive gel. A reference electrode was placed on the participant’s left clavicle. EMG signals were amplified (gain = 10^3^), and then digitally sampled at 1,000 Hz. EMG data were then band-pass filtered (20–500 Hz, 2-pass, 2nd-order Butterworth) and full-wave rectified.

In Experiments 1, 2, and 3 we investigated whether shoulder muscle activity adapted after learning to elbow reaches with shoulder fixation in a distinct shoulder orientation, distinct elbow orientation and distinct reach distance/speed. To compare the changes in the amplitude of muscle activity over trials and across different phases of the protocol, we calculated the mean amplitude of agonist muscle activity across a fixed time-window, −100 ms to +100 ms relative to movement onset, as has been done previously (see Debicki and Gribble 2005; Maeda et al. 2017, 2018).

In Experiment 4, we investigated whether feedback responses of shoulder muscles in a distinct shoulder orientation also adapt following learning of novel limb dynamics with shoulder fixation. To test whether the short and long latency stretch response of shoulder extensor account for and adapt to novel intersegmental dynamics, we binned the PEC EMG into previously defined epochs (see Pruszynski et al. 2008). This included a pre-perturbation epoch (PRE, −50-0 ms relative to perturbation onset), the short-latency stretch response (R1, 25-50 ms), the long-latency stretch response (R2/3, 50-100 ms), and the voluntary response (VOL, 100-150ms).

Normalization trials prior to each experiment were used to normalize muscle activity such that a value of 1 represents a given muscle sample’s mean activity when countering a constant 1 Nm torque (see Maeda et al. 2017, 2018; Pruszynski et al. 2008). Data processing was performed using MATLAB (r2017b, Mathworks, Natick, MA).

### Statistical analysis

Statistical analyses were performed using R (v 3.2.1). We performed different statistical tests (e.g., repeated measures ANOVA with Tukey tests for multiple comparisons, and t-tests), when appropriate in each of the experiments. Details of these analyses are provided in the Results section. Experimental results were considered statistically significant if the corrected p-value was less than < 0.05.

## Results

### All Experiments: Reducing shoulder muscle activity following shoulder fixation

In all experiments, participants moved their hand-cursor from a home target to a goal target placed along an arc such that it could be achieved by rotating only the elbow joint. We did not enforce a particular trajectory, but participants still performed the task by almost exclusively rotating only their elbow joint (Maeda et al. 2017, 2018).

To accomplish this movement, participants needed to compensate for the torques that arise at the shoulder when the forearm rotates around the elbow. Consistent with this requirement, in all experiments and arm postural conditions, we found substantial shoulder extensor muscle activity prior to movement onset (Gribble and Ostry 1999; Maeda et al. 2017); (Figures 2-5). After these baseline trials, we locked the shoulder joint of the robot with a physical clamp attached to the robotic apparatus and participants were then required to continue generating elbow extension movements (reach learn). Locking the shoulder joint eliminates the interaction torques that arise at the shoulder when the forearm rotates around the elbow and thus removes the need to activate shoulder muscles. Note that this manipulation also clamps reaching trajectories and did not alter task performance, with participants continuing to demonstrate >90% success rates.

**Figure 5:**
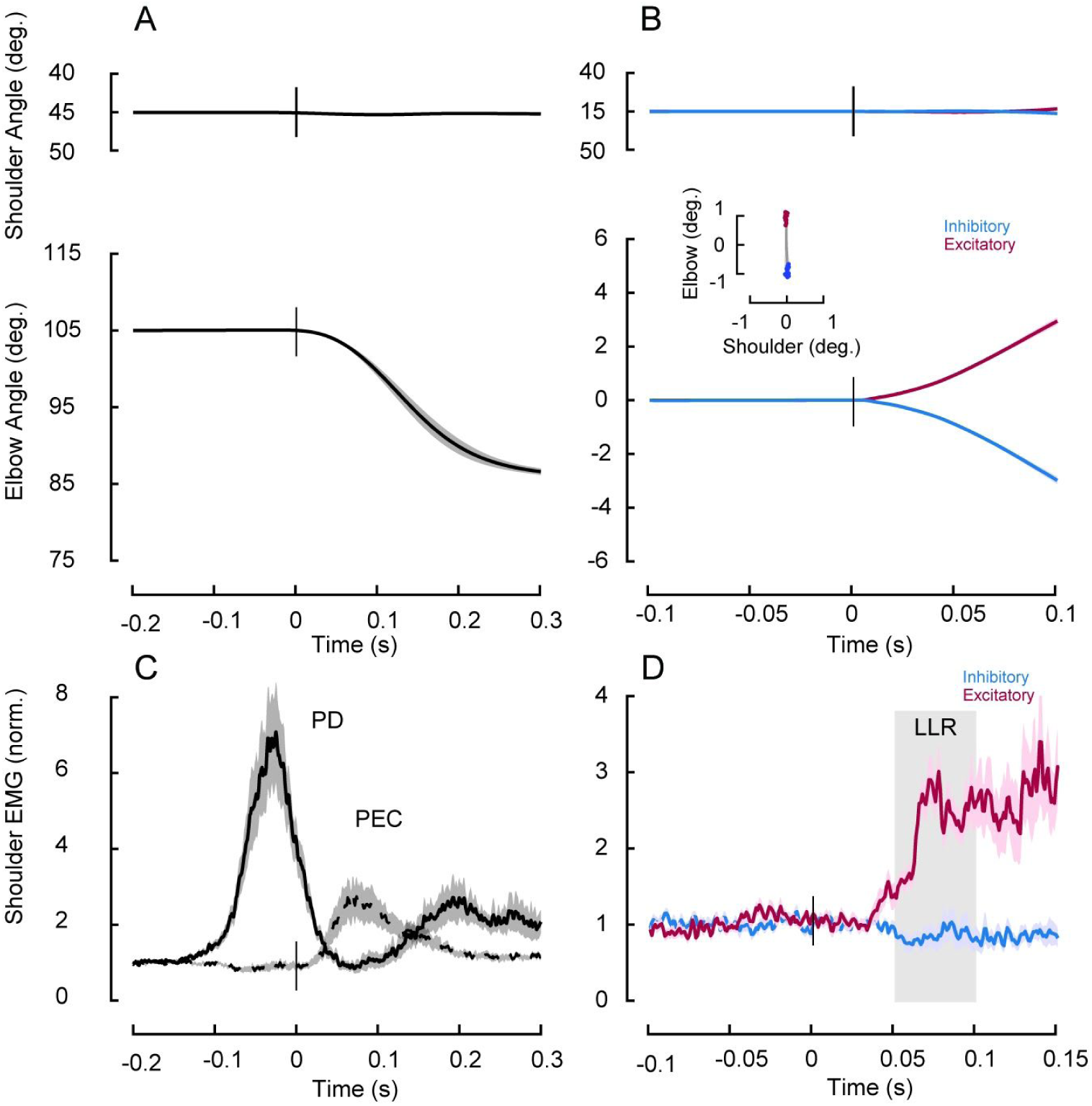
Baseline reaching and perturbation behaviour in Experiment 4. **A**, Average shoulder and elbow angle of elbow extension trials. Shaded error areas represent the standard error of the mean (SE). Data are aligned on movement onset. **B**, Average shoulder and elbow angle following mechanical perturbations that created pure elbow motion. Red and blue traces are from the shoulder/elbow extensor torque and shoulder/elbow flexor torque conditions, respectively. Shaded error areas represent the SE. Data are aligned on perturbation onset. Inset shows the amount of shoulder and elbow displacement 50 ms post-perturbation (data are shown for all subjects). **C**, Solid and dashed lines represent average agonist (PD) and antagonist (PEC) muscle activity during elbow extension movements. EMG normalized as described in the Methods. Data aligned on movement onset. Shaded error areas represent the standard error (SE). **D**, Normalized shoulder (PD) muscle activity associated with panel **B**. Data aligned on perturbation onset. Shaded error areas represent the SE. Gray horizontal shaded areas represent the long-latency reflex epoch (LLR).

We first confirmed that the nervous system reduces shoulder muscle activity when the shoulder is locked (Maeda et al. 2018). Figure 6 A-B shows the mean shoulder extensor muscle activity in a fixed time window (−100 to 100ms centered on movement onset, see Methods) over elbow extension movements before locking the shoulder joint (i.e. baseline phase), with the shoulder locked (i.e adaptation phase) and after unlocking the shoulder joint again (i.e. post-adaptation phase). Consistent with our previous work, we found that the magnitude of shoulder muscle activity slowly decayed over the adaptation trials and quickly returned to baseline levels after removing the shoulder lock in the post-adaptation phase (Figure 6 A-B). We performed one-way ANOVAs for each experiment to compare shoulder agonist muscle activity across learning phases (last 25 trials in the baseline trials, adaptation and post-adaptations phases, see Maeda et al., (2018)). We found a reliable effect of phase in each experiment (Experiment 1: F_2,28_ = 13.53, P < 0.0001; Experiment 2: F_2,28_ = 23.05, P < 0.0001; Experiment 3: F_2,28_ = 7.971, P = 0.0018; Experiment 4: F_2,28_ = 4.05, P = 0.01). Tukey post-hoc tests showed a reliable reduction of shoulder muscle activity from baseline to adaptation phases in each experiment (Experiment 1: 26.4%, P < 0.0001; Experiment 2: 30%; P < 0.001; Experiment 3: 25.8%; P < 0.001; Experiment 4: 23.2%; P = 0.01).

**Figure 6:**
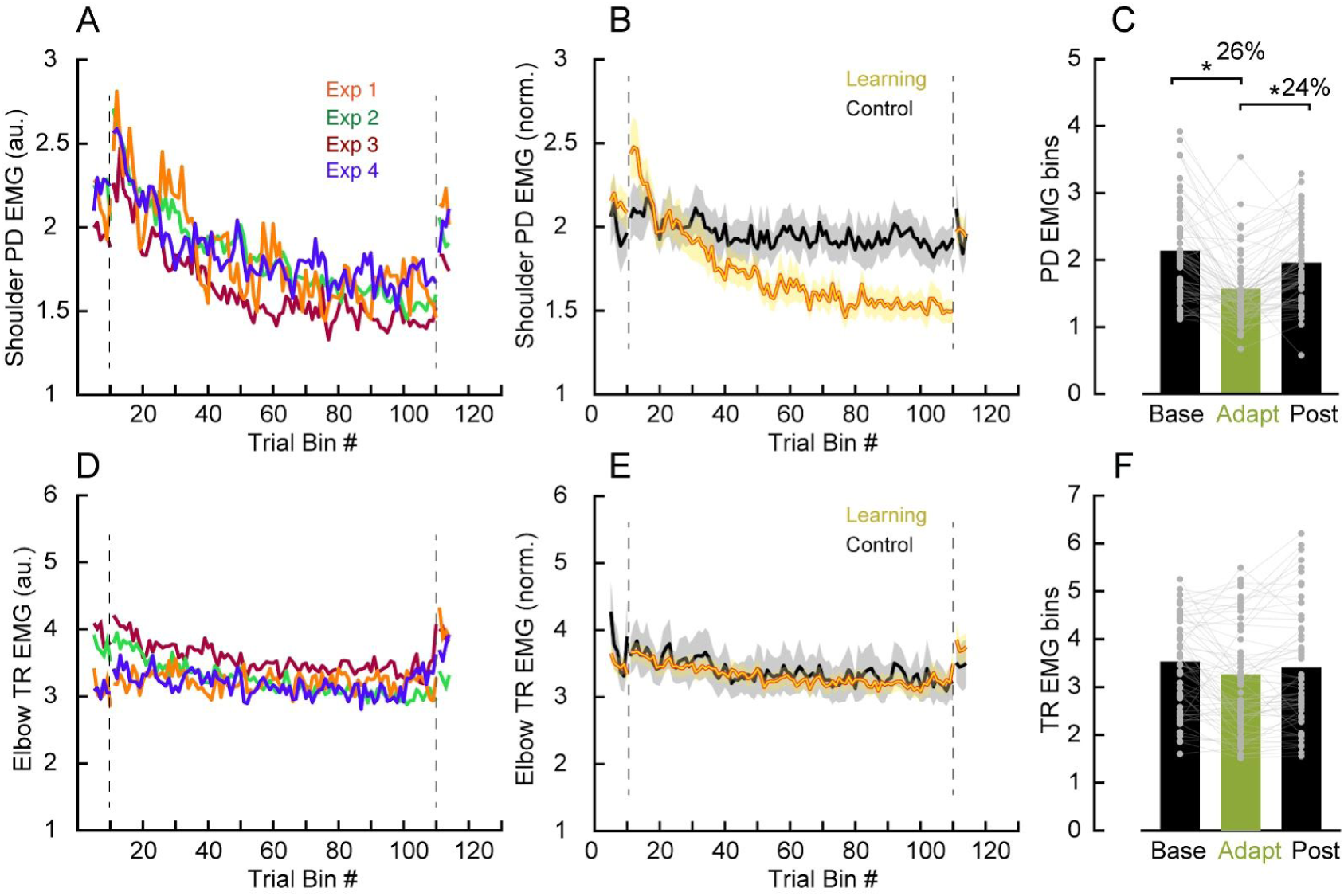
Reducing shoulder muscle activity following shoulder fixation. **A**, Average PD muscle activity in a fixed time window (−100 to 100 ms relative to movement onset). Each data bin is 5 trials. EMG normalized as described in Methods. Traces represent data from each of the 4 experiments. **B**, Average PD muscle activity in a fixed time window. Each data bin is 5 trials. Traces represent data combined across Experiments 1-4. Black traces are from the control experiment. Shaded error areas represent the standard error (SE). **C**, Average of the last 5 bins of trials in each phase of the protocol. Each dot represents data from a single participant. Asterisks indicate reliable effects (p<0.05, see main text). **D-F**, Data for elbow TR muscle in same format as **A-B**.

We then merged the data from all experiments and performed a one-way ANOVA to compare shoulder agonist muscle activity across learning phases in all experiments. Again, we found an effect of phase (F_2,156_ = 31.43, P < 0.0001) on shoulder muscle activity. Tukey post-hoc tests showed that shoulder extensor muscle activity decreased by 26% relative to baseline (P < 0.0001) with the shoulder locked and increased again after unlocking the shoulder joint, quickly returning to baseline trials (P = 0.17; Figure 6 A-C). We found no reliable changes in monoarticular elbow TR muscle as a function of phases (one-way-ANOVA, F_2,156_ = 1.56, P = 0.193) and experiment (two-way-ANOVA, F_2,156_ = 0.68, P = 0.5; Figure 6 D-F).

We also performed a control experiment in which participants (N=10) performed the same task with the same number of trials but never experienced the shoulder lock manipulation. We found no corresponding effect of phase (One-way ANOVA, F_2,18_ = 0.579, P = 0.571) on shoulder agonist muscle activity (Figure 6 B, see black traces). This result rules out the possibility that the decay of shoulder muscle activity relates to extensive experience performing elbow rotations rather than our shoulder fixation manipulation.

Lastly, we performed a second control experiment (see Methods) in which participants (N = 30) performed the same elbow reaching task and adapted to shoulder fixation (i.e. baseline – shoulder unlocked, adaptation – shoulder locked and post-adaptation phase – shoulder unlocked) in the generalization conditions of Experiments 1 (distinct shoulder initial orientation, as in (Maeda et al. 2018), 2 (distinct elbow initial orientation) and 3 (distinct distance/speed). This control experiment was performed to confirm that participants reduce shoulder muscle activity with shoulder fixation when the training takes place in the generalization conditions (Maeda et al. 2018). We found a reliable effect of phase in each experiment (Experiment 1: F_2,18_ = 7.585, P = 0.004; Experiment 2: F_2,18_ = 6.34, P = 0.008; Experiment 3: F_2,18_ = 7.77, P = 0.003). Tukey post-hoc tests showed a reliable reduction of shoulder muscle activity from baseline to adaptation phases in each experiment (Experiment 1: 33.6%, P = 0.001; Experiment 2: 42.8%; P = 0.005; Experiment 3: 21.06%; P < 0.009).

### Experiment 1: Generalization to reaching in a different initial shoulder configuration

In this experiment, we tested whether learning new intersegmental dynamics following shoulder fixation during a 25 degree elbow extension movement starting at one initial shoulder orientation (reach learn) generalizes to the same 25 degree elbow extension movement but starting with a different initial shoulder orientation (reach generalize) (Figure 2 A-D, baseline trials). Participants (N=15) had no difficulty performing the task with the imposed speed and accuracy constraints at either configuration and did so with >90% success within 5 minutes of practice.

We used a one-way ANOVA to compare shoulder extensor muscle activity in the baseline trials, late in the adaptation trials and late in the post-adaptation trials in the generalization posture (reach generalize), and we found a reliable effect of phase on shoulder activity (F_2,28_ =11.03, P < 0.0001) (Figure 7 A-B). Tukey post-hoc tests showed that, in the generalization condition, shoulder extensor muscle activity decreased by 24% relative to baseline (P < 0.001) with the shoulder locked. To contrast this reduction in shoulder activity in the generalization posture (reach generalize) with respect to the reduction in shoulder muscle activity in the learning posture (reach learn), we performed a paired t-test between the difference in shoulder muscle activity in the learning and generalization postures with respect to their respective baseline (last 10 trials). We found similar reduction of shoulder muscle activity in the generalization posture (reach generalize) compared to the shoulder activity in the learned posture (reach learn; t_14_ = −0.43, P = 0.67; Figure 7 A inset), suggesting that this generalization was complete.

**Figure 7:**
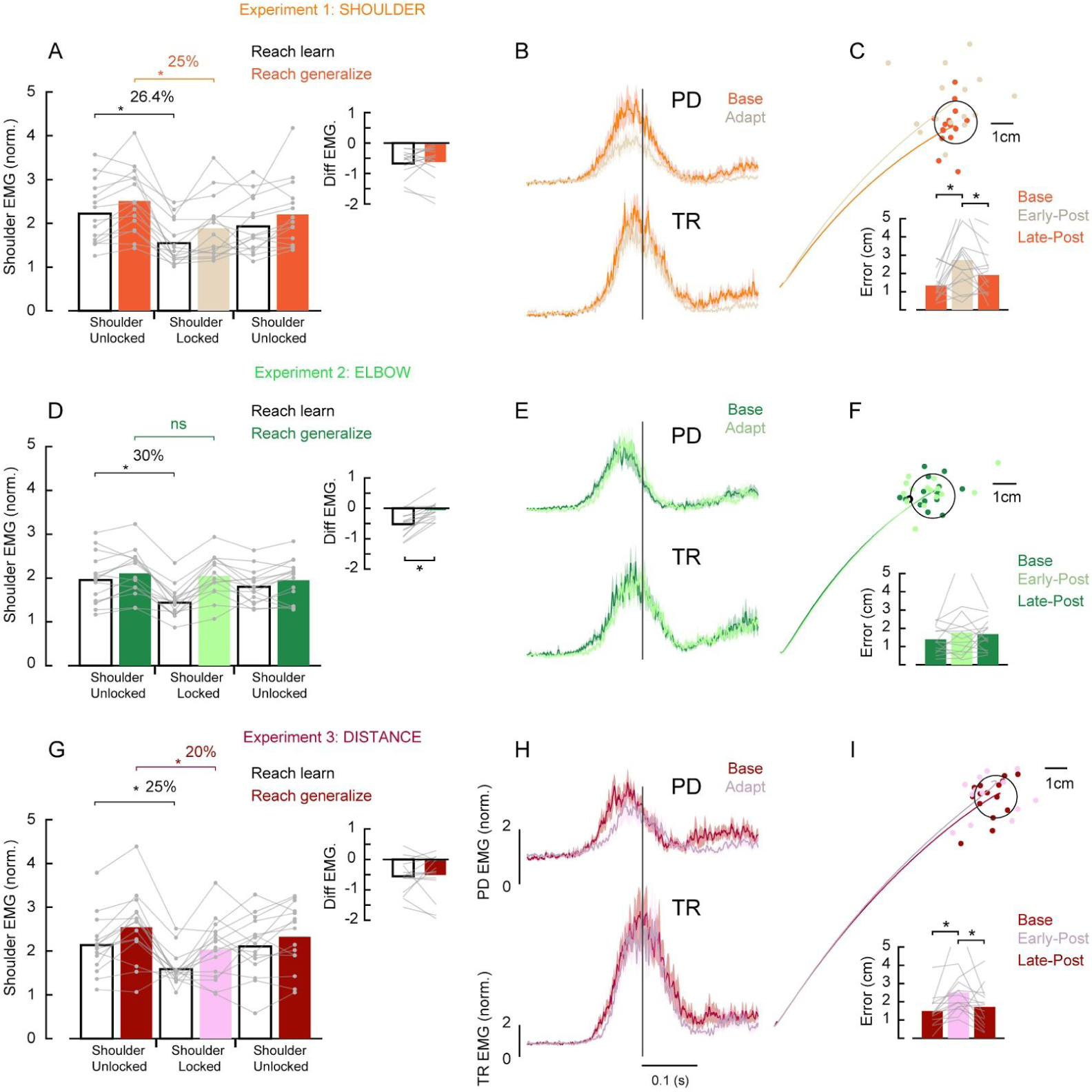
Generalization of learning of novel intersegmental dynamics to distinct elbow reaches. **A**, Average PD muscle activity in a fixed time window (−100 to 100 ms relative to movement onset) of reaches in the baseline (10 trials), at the end of adaptation (10 trials) and early in the post-adaptation (10 trials) phases in the two different initial shoulder orientation (reach learn and reach generalize, experiment 1). Inset shows the difference in shoulder muscle activity (the end of adaptation - baseline phases) for the two postures. Each line represents an individual participant. **B**, Time series of shoulder PD (upper panel) and elbow TR (lower panel) muscle activity in the baseline (10 trials), and the end of adaptation (10 trials) in the generalization orientation. Shaded error areas represent the standard error of the mean (SE). Data are aligned on movement onset. **C**, Average hand trajectories late in the baseline (10 trials) and early in the post-adaptation trials (first 3 trials) associated with experiment 1. Inset bar graph shows the average error between hand position at movement offset to the center of the target in the last 10 trials in the baseline, first 3 trials early in the post-adaptation and last 10 trials late in post-adaptation phases. Each dot represent data from a single participant. **D-F**, Data associated with the experiment starting in a different initial elbow orientation (reach learn and reach generalize, experiment 2). Data in the same format as A-C. **G-I**, Data associated with the experiment starting in a different reach distance/speed (reach learn and reach generalize, experiment 3). Data in the same format as A-C. Asterisks indicate reliable effects (p<0.05, see main text).

We also tested whether there was evidence of kinematic aftereffects in the direction predicted when failing to compensate for the now-unlocked shoulder joint (Figure 7 C) (Maeda et al. 2018). We performed a one-way ANOVA to compare reach kinematics (at 80% of the movement, see Methods) of trials late in the baseline phase (last 25 trials) with trials early in the post-adaptation phase (first 3 trials) (Figure 7 C). We assessed only the first three trials after unlocking the shoulder joint because the aftereffect washes out very quickly. We found an effect of phase on kinematic traces (F_2,28_ = 7.2, P = 0.002). Tukey post-hoc tests showed that kinematic traces early in the post adaptation increased relative to baseline (P = 0.001) and returned to baseline levels in late post-adaptation (P = 0.25; Figure 7 C).

### Experiment 2: Generalization to reaching in a different initial elbow configuration

In this experiment, we tested whether learning new intersegmental dynamics following shoulder fixation during a 25 degree elbow extension movement starting at one initial elbow orientation (reach learn) generalizes to a 25 degree elbow extension movement starting at a different initial elbow orientation (reach generalize) (Figure 3 A-D, baseline trials). Participants (N=15) had no difficulty performing the task with the imposed speed and accuracy constraints and did so with >90% success in both configurations within 5 minutes of practice.

We again used a one-way ANOVA to compare shoulder extensor muscle activity in the baseline trials, late in the adaptation and late in the post-adaptation trials in the generalization posture. Here we found no effect of phase on shoulder activity indicating a lack of generalization (F_2,28_ = 2.43, P = 0.1) (Figure 7 D-E). As expected, given this negative result, a paired t-test between the difference in shoulder muscle activity in the learning and generalization postures with respect to their respective baseline (10 trials) showed a reliable effect, indicating that shoulder muscle activity in the generalization posture did not follow the reduction observed in the learning posture (t_14_ = −4.97, P < 0.001; Figure 7 D inset).

We also performed a one-way ANOVA to compare reach kinematics (at 80% of the movement) of trials late in the baseline phase (last 25 trials), and trials early in the post-adaptation phase (first 3 trials) and trials late in the post-adaptation phase (last 25 trials). Consistent with the lack of shoulder muscle activity changes, we found no effect of phase on reach trajectories (F_2,28_ = 0.92, P = 0.41, Figure 7 F).

### Experiment 3: Generalization to a different reach distance/speed

In this experiment, we tested whether learning new intersegmental dynamics during pure elbow rotation with shoulder fixation generalizes to larger elbow rotations starting from the same initial position. That is, participants were required to make either 20° (reach learn) or 30° (reach generalize) elbow extensions from the same initial configuration (Figure 4 A-D, baseline trials). Participants (N=15) had no difficulty achieving the speed and accuracy constraints for either reach condition with >90% success within 5 minutes of practice.

We used a one-way ANOVA to compare shoulder extensor muscle activity in the baseline trials, late in the adaptation trials and late in the post-adaptation trials in the generalization posture, and we found a reliable effect of phase on shoulder activity (F_2,28_ = 6.194, P = 0.005). Tukey post-hoc tests showed that shoulder extensor muscle activity decreased by 20% relative to baseline (P < 0.001) with the shoulder locked and increased again after unlocking the shoulder joint, returning to baseline trials (Figure 7 G-H). We also found that shoulder muscle activity in the two reaching conditions late in the adaptation phase (difference baseline - adaptation) were not reliably different, suggesting that the generalization that took place was complete with respect to the amount of shoulder muscle activity reduction exhibited in the learning condition (t_14_ =-0.26, P = 0.79; Figure 7 G inset).

Lastly, we again investigated whether there was an evidence of reliable after-effects. To do so we performed a one-way ANOVA to compare reach kinematics (at 80% of the movement) of trials late in the baseline phase (last 25 trials), and trials early in the post-adaptation phase (first 3 trials) and trials late in the post-adaptation phase (last 25 trials). We found an effect of phase on this kinematic variable (F_2,28_ = 4.9, P = 0.01, Figure 7 I). Tukey post-hoc tests showed that kinematics deviated early in the post-adaptation phase relative to baseline (P = 0.007) but returned to baseline levels late in the post-adaptation trials (P = 0.77).

### Experiment 4: Generalization to feedback responses in a different initial shoulder configuration

The main goal of Experiment 4 was to examine whether learning novel intersegmental dynamics following shoulder fixation during feedforward (i.e. voluntary—reach learn) control modifies the sensitivity of feedback (i.e. reflex generalize) responses to mechanical perturbations applied in a different shoulder orientation (Figure 5 A-B, baseline trials). The mechanical perturbations consisted of 100 ms step torques applied concomitantly to the shoulder and elbow such that it caused minimal shoulder motion but distinct amounts of elbow motion (Figure 5, inset). Participants (N=15) had no difficulty performing the task and did so with >90% success within 5 minutes of practice.

As previously demonstrated, participants generated a substantial amount of shoulder muscle activity during reaching trials (Gribble and Ostry 1999; Maeda et al. 2017, 2018). Also, as previously demonstrated, we found that mechanical perturbations that created pure elbow motion elicited substantial shoulder muscle activity in the long-latency epoch (Figure 5 D), as appropriate for countering the imposed joint torques (Kurtzer et al. 2008; Maeda et al. 2017, 2018).

Figure 8 illustrates the mean difference of shoulder (PD) muscle activity between excitatory and inhibitory torque perturbation trials. Traces are shown before and after learning the novel intersegmental dynamics during reaching. We performed a one-way ANOVA to compare the difference of PD muscle activity in the long-latency epoch across trials in the baseline, adaptation and post-adaptation phases. We found a reliable effect of phase (F_2,28_ = 6.61, P = 0.004). Tukey post-hoc tests showed that the difference in PD muscle activity in the long-latency epoch decreased by 34% (P = 0.001) following shoulder fixation and returned to baseline levels in the post-adaptation phase (P = 0.82) (Figure 8 A-B). We performed the same analysis to test for changes in the short-latency epoch, but we found no reliable differences (F_2,28_ = 0.061, P = 0.9). We also tested whether there was a change in baseline EMG activity pre-perturbation across phases, which could explain these changes in EMG in the long-latency epoch (gain scaling, (Pruszynski et al. 2009). We used a one-way ANOVA to compare the baseline activity of PD muscle activity pre-perturbation as a function of experimental phase but also found no reliable effect (F_2,28_ = 0.712, P = 0.49; Figure 5 B, inset). We also found no corresponding changes in both long-latency (one-way-ANOVA, F_2,28_ = 1.75, P = 0.19) and short-latency epochs (one-way-ANOVA, F_2,28_ = 0.663, P = 0.52) of the monoarticular elbow TR muscle. There was also no reliable change in the baseline activity of TR muscle activity pre-perturbation as a function of phases (F_2,28_ = 1.26, P = 0.29).

**Figure 8:**
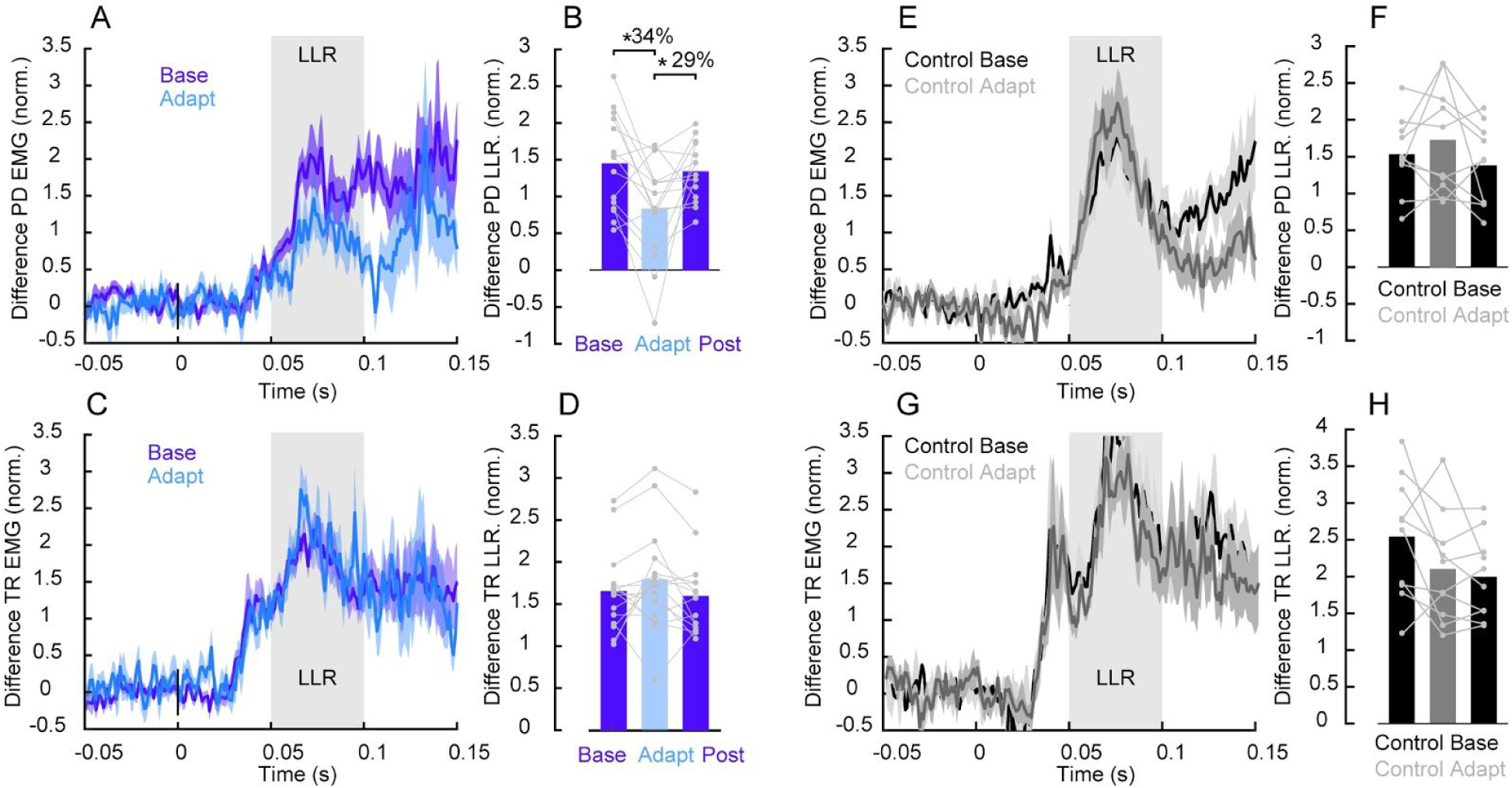
Rapid feedback responses in a different shoulder initial orientation. **A**, Average shoulder PD EMG. (filtered, rectified and normalized as described in Methods). Data aligned on perturbation onset. Shaded error areas represent the SE. Dark and light blue traces are trials in the baseline and after learning, respectively. **B**, Average shoulder PD EMG. of the long latency epoch (50-100ms) across trials late in the baseline, after learning and late in the post-adaptation phases. Each dot represent data from a single participant. Asterisks indicate reliable effects (p<0.05, see main text). **C-D**, Data for elbow TR muscle in same format as **A-B**. Gray horizontal shaded areas represent the long-latency reflex epoch (LLR). **E-H**, Data for the control study in which the shoulder joint was never locked shown in the same format as **A-D.**

In a control experiment, participants (N=10) performed the same task with the same number of trials but never experienced the shoulder lock manipulation. We found no corresponding effect of phase (one-way ANOVA, F_2,18_ = 2, P = 0.16) on shoulder PD in the long-latency epoch (Figure 8 E-F).

## Discussion

In Experiment 1, we tested whether learning to reduce shoulder muscle activity for pure elbow rotations after shoulder fixation generalizes to pure elbow rotations in a new initial shoulder orientation. Consistent with generalization, we found reduced shoulder muscle activity in the new shoulder orientation. In Experiment 2, we tested whether learning to reduce shoulder muscle activity for pure elbow rotations after shoulder fixation generalizes to pure elbow rotations in a new initial elbow orientation. We found no evidence of generalization. Shoulder muscle activity in the new elbow orientation did not decrease relative to levels without fixation at that new orientation. In Experiment 3, we tested whether learning to reduce shoulder muscle activity for pure elbow rotations after shoulder fixation generalizes to larger amplitude/speed of elbow rotations in the same arm configuration. Consistent with generalization, we found reduced shoulder muscle activity for larger amplitude/speed of rotations compared to levels without fixation for the larger amplitude/speed. In Experiment 4, we tested whether learning to reduce feedforward (i.e. agonist) shoulder muscle activity for pure elbow rotations after shoulder fixation generalizes to feedback (i.e. reflex) responses following mechanical perturbations in a new shoulder orientation. Consistent with generalization and transfer, we found a reduction in shoulder feedback responses in the new shoulder orientation.

### Generalization across the workspace

We have previously demonstrated that participants learn to decrease shoulder muscle activity when performing elbow flexion and extension movements following shoulder fixation (Maeda et al. 2018). This manipulation alters normal arm dynamics by eliminating the interaction torques that arise at the shoulder due to forearm rotation. Here we investigated how this type of learning is represented by looking at whether and how it generalizes to movements made in different joint configurations and thus parts of the workspace (Krakauer et al. 2019; Shadmehr 2004). Given that this type of learning unfolds slowly, in the absence of kinematic errors and that it is incomplete, we previously suggested that the nervous system may attribute our shoulder fixation manipulation to a physical change of the body as opposed to a changing environment or the feature of some hand-held tool (Maeda et al. 2018). If this were the case, then such learning should generalize broadly, presumably to all different joint orientations and speeds/distances. However, this does not appear to be the case as, consistent with previous theoretical and empirical work investigating generalization in other learning contexts, such as force field and visuomotor learning, we found a mixed generalization pattern (Berniker et al. 2014; Brayanov et al. 2012; Burgess et al. 2007; Criscimagna-Hemminger et al. 2003; Donchin et al. 2003; Gonzalez Castro et al. 2011; Goodbody and Wolpert 1998; Joiner et al. 2011; Krakauer et al. 2000, 2006; Malfait et al. 2002, 2005; Mattar and Ostry 2007, 2010; Pearson et al. 2010; Shadmehr and Moussavi 2000; Thoroughman and Shadmehr 2000)

Some insight into the generalization pattern can be gained by examining the joint torques required to generate pure elbow motion at the learning and generalization configurations of Experiments 1-3 (see insets in Figures 2-4). In Experiment 1, changing initial shoulder orientation does not change the underlying shoulder and elbow torques required to produce pure elbow rotation. Thus, any change in shoulder muscle activity learned for one configuration is directly applicable to the new joint configuration. The fact that we found generalization is therefore not particularly surprising but rules out the possibility that this learning is spatially limited to some local part of the workspace. Both Experiments 2 and 3 do require changing shoulder and elbow torque but we only observe generalization in the latter. Changing the elbow orientation, as in Experiment 2, requires changing how the shoulder and elbow joints work together to achieve pure elbow rotation. In the configurations we used, pure elbow rotation in the generalization configuration requires less elbow torque but more shoulder torque. Increasing the distance and speed from the same initial orientation, as in Experiment 3, does not change how the two joints need to be coordinated. That is, executing a larger and faster movement requires simultaneous scaling of both shoulder and elbow torque and, in the configurations we used, this scaling factor was very similar across the two joints. The pattern we report is reminiscent of previous studies examining learning visuomotor distortions where learning a new gain, which produce larger errors, broadly generalize across the workspace whereas learning rotations which produce directional errors was limited (Krakauer et al. 2000). Taken together, our results suggest that the nervous system is not learning a new internal model of the arm’s dynamics but is consistent with the notion that the nervous system is implementing a habitual control scheme that is able to scale set coordination patterns but not change them (Debicki and Gribble 2004; de Rugy et al. 2012).

Why did the nervous not learn a new internal model of the arm’s dynamics? Two possibilities are that participants experienced altered arm dynamics (1) in a very limited context and (2) for a relatively short duration. In our study, participants only experienced shoulder fixation in a very small part of the workspace (one reach condition and only in the horizontal plane). Learning was slow with shoulder muscle activity showing a ∼30% reduction over ∼1000 trials performed over 1-2 hours. it remains unclear, however, whether extended practice in this protocol would be sufficient for people to overcome the massive experience they have with their normal limb. That is, there may exist a relatively synergy between shoulder and elbow muscles that is resistant to change even on the timescales of weeks (Debicki and Gribble 2004; de Rugy et al. 2012). These findings and ideas are also relevant for understanding recovery following movement impairments (long-term injuries, stroke) (Krakauer 2006) and for the development of rehabilitation approaches, such as those focusing on constraint-induced movement therapy, which has been shown useful for stroke patients by combining restraint of the unaffected limb and intensive practice with the affected limb (Dromerick et al. 2000; Krakauer 2006; Mark and Taub 2004).

Another key feature of our paradigm is that participants do not make substantial kinematic errors when the shoulder is fixed. Given that generalization of motor learning is often studied in movement tasks in which movement errors are present (ie. force field learning) (Krakauer et al. 2019) and that there is evidence that generalization happens as a function of the nervous system identifying the source of errors (Berniker and Kording 2008), it was not clear whether the nervous system would generalize following shoulder fixation. Here we found that learning generalized despite the task not introducing explicit errors indicating that such errors are not required to drive generalization and posing the question about whether these types of learning share underlying neural circuits (Maeda et al. 2018; Vaswani and Shadmehr 2013).

### Generalization and transfer to feedback control

Previous research has demonstrated that the sensitivity of feedback responses change over the course of motor learning (Ahmadi-Pajouh et al. 2012; Cluff and Scott 2013; Maeda et al. 2018; Wagner and Smith 2008). For instance, when participants learned to reach in the presence of force fields and encounter mechanical perturbations occasionally over the course of learning their feedback responses, 50 ms following perturbation onset (ie. long-latency epoch), are modified in a similar rate and direction observed during learning. In addition, we have previously demonstrated that these feedback responses also change when people learn a new intersegmental dynamics during elbow reaching with shoulder fixation (Maeda et al. 2018). In particular, we found a gradual decrease of shoulder muscle activity with shoulder fixation during reaching and a gradual decrease in these rapid feedback responses probed over the course of learning. It is currently unknown whether feedback responses also have access to subsequent features of motor learning such as generalization of learning. In our Experiment 1 and 4, we found that the nervous system generalizes similarly for feedforward control in a distinct shoulder orientation and to feedback control when perturbations were applied in a distinct shoulder orientation, respectively. These similar generalization patterns are consistent with the idea that feedforward and feedback control share an internal model for motor control (Ahmadi-Pajouh et al. 2012; Cluff and Scott 2013; Maeda et al. 2018; Wagner and Smith 2008). An interesting question from this view for future studies is whether these feedback responses would also have access to motor memories across sessions and days, such as savings (Krakauer 2009; Krakauer et al. 2019).

A direct investigation of the neural mechanisms that allow the nervous system to generalize by scale a set of coordination patterns for feedforward and feedback control is a largely unexplored topic. The prediction is that such scaling for this learning and generalization should happen in overlapping neural circuits for feedforward and feedback control. One potential area of interest is the primary motor cortex (M1). Gritsenko et al. (2011) applied transcranial magnetic stimulation to human M1 while participants reached to targets displayed around the workspace such that the upcoming movement yielded assistive or resistive interaction torques between the shoulder and elbow joints. They found greater motor evoked potentials for reaching under the resistive interaction torques compared to assistive, which indicates that M1 mediates feedforward control of intersegmental dynamics. Pruszynski et al. (2011) also applied transcranial magnetic stimulation to human M1 and found that it potentiates shoulder muscle responses following mechanical perturbations that cause pure elbow motion, indicating that M1 also mediates feedback control of intersegmental dynamics. There is also evidence from a single neuron level that neurons in M1 are activated during reaching also respond to mechanical perturbations (Evarts 1973; Evarts and Fromm 1981; Evarts and Tanji 1976; Herter et al. 2009; Omrani et al. 2014; Picard and Smith 1992; Pruszynski et al. 2011, 2014; Wolpaw 1980).

Another potential region of interest is the cerebellum. Research has demonstrated that patients with damage in the cerebellum show deficits in coordinating joints without affecting the ability to generate forces (Bastian et al. 1996, 2000; Goodkin et al. 1993; Holmes 1939). Long-latency feedback responses following mechanical perturbations that create pure elbow motion have been reported to be reduced in patients with cerebellar dysfunction, consistent with the view that this region also mediates feedback control of intersegmental dynamics (Kurtzer et al. 2013). Interestingly, research has also considered the cerebellum for hosting multiple paired internal models (Wolpert et al. 1998). This could be relevant in the context of our finding that learning generalizes to some conditions but not in others, as it may involve different paired internal models rather than a single one. Alternatively, an internal model in the cerebellum has also been also viewed from the perspective that it is composed with elements that dictate generalization patterns (Shadmehr 2004). An important topic for future research is to identify whether and how these set of elements that support generalization are present in the neural code for feedforward and feedback control and the interaction between areas, including M1 and cerebellum that might support its implementations.

## Notes

Grants: This work was supported by a grant from the National Science and Engineering Research Council of Canada (NSERC Discovery Grant to J.A.P.: RGPIN-2015-06714). R.S.M. received a salary award from CNPq/Brazil. J.M.Z. received an Undergraduate Student Research Award from the National Science and Engineering Research Council of Canada. J.A.P. received a salary award from the Canada Research Chairs program.

#### Summary of Updates

Added control experiments to ensure learning can occur in the generalization postures. Several other changes in presentation of the material and associated analyses.

